# Video-rate remote refocusing through continuous oscillation of a membrane deformable mirror

**DOI:** 10.1101/2021.04.27.441576

**Authors:** Terry Wright, Hugh Sparks, Carl Paterson, Chris Dunsby

## Abstract

This paper presents the use of a deformable mirror (DM) configured to rapidly refocus a microscope employing a high numerical aperture objective lens. An Alpao DM97-15 membrane DM was used to refocus a 40×/0.80 NA water-immersion objective through a defocus range of −50 to 50 μm at 26.3 sweeps per second. We achieved imaging with a mean Strehl metric of > 0.6 over a field of view in the sample of 200×200 μm^2^ over a defocus range of 77 μm. We describe an optimisation procedure where the mirror is swept continuously in order to avoid known problems of hysteresis associated with the membrane DM employed. This work demonstrates that a DM-based refocusing system could in the future be used in light-sheet fluorescence microscopes to achieve video-rate volumetric imaging.

## 1. Introduction

The ability to refocus an optical microscope rapidly, on sub-second timescales is essential for example when using optically-sectioning microscopy to acquire 3D images of the specimen at multiple volumes per second [1]. The most straightforward approach to achieve refocusing is axial translation of the sample or microscope objective with a piezoelectric actuator [2]. However, moving the mass of the sample or lens at the required frequency and amplitude needs an actuator of relatively high power and can introduce vibrations that can perturb the sample.

An alternative to moving the sample or lens is to employ an adjustable optical element elsewhere in the optical system. Electrically tunable lenses (ETL) provide convenient electronic control of the amount of defocus [3] and have been successfully demonstrated to achieve remote refocusing for multiphoton microscopy [4], light-sheet microscopy [5] and confocal microscopy [6]. The ETL in [5] was able to scan 17 planes within a zebrafish heart at 30 volumes per second. The authors used a medium/low NA lens which corrected for primary defocus and they reported that astigmatism, coma and field curvature restricted the use of the lens to the central region of the field of view: these aberrations increased towards the limits of the ETL focal range. ETLs in general only provide low-order quadratic primary defocus wavefront correction whereas spherical aberration and higher orders are required to remotely refocus high-NA objectives [7]. The proposed bi-actuator design of a liquid ETL in [8] consisting of 2 concentric piezo rings was able to provide simultaneous correction for defocus and spherical aberration. However, more degrees of freedom are required for greater aberration correction of high-NA lenses and when imaging in non-homogeneous media. ETLs exhibit significant hysteresis; a study of ETLs produced by Optotune [9] reported hysteresis up to 1D of optical power that was dependent on the history of the applied current used to refocus the lens.

We will refer to the total wavefront correction required as high-NA defocus, see equation 1 in reference [7]. Figure 1 plots the maximum diffraction-limited refocus that can be achieved as a function of NA when applying only quadratic defocus wavefront correction (see Appendix for calculation). For an NA of 0.75 the application of primary defocus alone only allows a diffraction-limited remote refocus distance of 22 μm with a water immersion objective, and only 7 μm with an air objective giving full ranges of 44 μm and 14 μm respectively.

**Figure 1.**
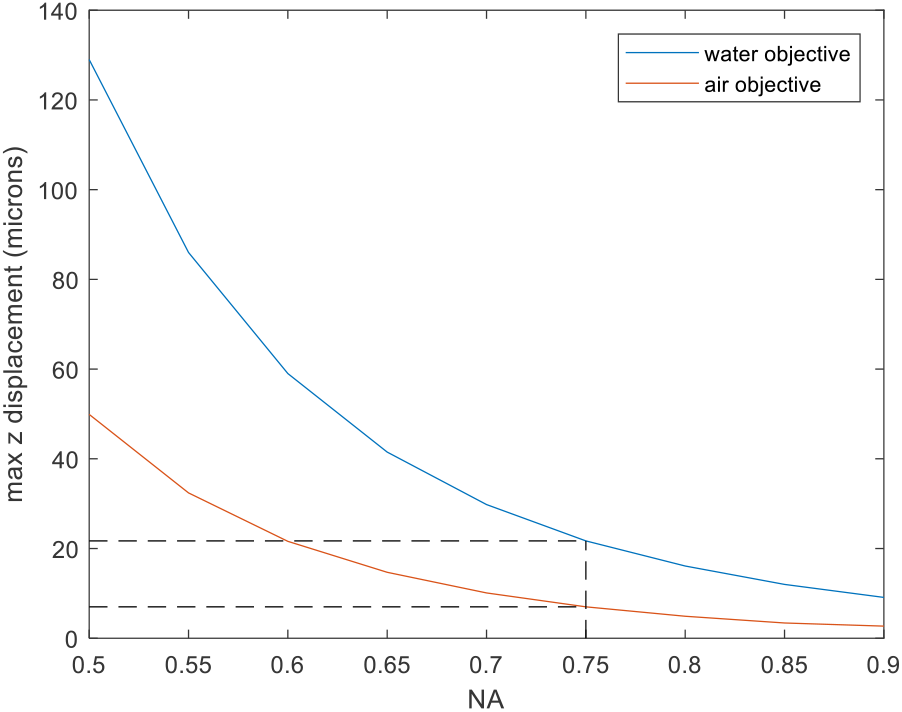
Maximum diffraction-limited refocus range (axial displacement) achievable for different NA with quadratic defocus correction. The dashed line indicates the maximum axial displacement for a system with NA=0.75 as used here. The blue curve is for a water immersion objective and the red is for an air objective.

In a two-photon fluorescence microscope a liquid crystal spatial light modulator (SLM) can be used to apply high-NA defocus to the excitation beam [12]. This approach is effective for two-photon imaging, which, due to the inherent sectioning afforded by two-photon excitation, does not require refocusing of the return signal, but SLM losses limit the application of this approach to other modalities.

The remote-refocusing approach of Botcherby et al. [7] can also achieve axial refocusing. Here, three microscopes are placed in series with an ideal intermediate image being formed between the 2^nd^ and 3^rd^ microscope objectives. Axial translation of the 2^nd^ or 3^rd^ microscope objectives achieves remote refocusing with no mechanical perturbation of the sample. However, in this configuration the actuator driving the translation of the 2^nd^ or 3^rd^ microscope objective still must be sufficiently powerful to accelerate the mass of the objective at the required frequency and amplitude. The mass being moved can be reduced by folding the optical system about the intermediate image, with remote refocusing achieved by scanning the axial position of the small fold mirror, but then a beam-splitter arrangement is required to separate the refocused beam. Remote refocusing of the excitation beam path can be performed without loss of light using a polarising beam splitter arrangement e.g. for scanning a polarised excitation spot in multiphoton microscopy [10], but if employed on the detection path it typically leads to a reduction in light throughput [11].

Rapid refocusing using a deformable mirror (DM) has been demonstrated previously for applying low-order defocus in optical coherence tomography [13]; for achieving an active focus lock in a confocal microscope [14]; and to extend the depth of field in an epi-fluorescence microscope [15]. Higher-order defocus wavefront correction with a DM has been demonstrated previously for rapid adaptive focusing in a multiphoton microscope [16]. The authors used open-loop control of an Alpao DM to acquire 10 optically-sectioned images axially separated by 5 μm in 10 ms from a fluorescently labelled pollen grain and 5 optically-sectioned images axially separated by 5 μm in a live fruit fly brain expressing a genetically encoded calcium indicator at 2 volumes/sec.

Higher-order membrane DMs additionally offer the possibility to correct for spherical, and higher-order aberrations, provide average corrections to some field-dependent aberrations such as field curvature, and compensate for system aberrations.

Changes in refractive index caused by local structures, e.g. proteins, nuclear acids and lipids have a refractive index different to water, can give rise to optical aberrations. Therefore, as well as refocusing the objective, another benefit of using DMs is that they can be used to correct for sample-induced aberrations, and this becomes more important when employing higher numerical aperture microscope lenses. For example, Débarre et al. [17] demonstrated an adaptive optics-based approach using a DM to correct for sample-induced aberrations when imaging to a depth of 140 μm in a mouse embryo. Sample-induced aberrations have been shown to be stable over hours when imaging in vivo in mouse brain [18], so once determined, aberrations can be corrected by a DM over a period of time-lapse imaging.

Alpao DMs are known to suffer from a viscoelastic creep on a timescale of minutes. This is suspected to be due to a polymer material used between the actuators and the mirror membrane surface, and so their response is dependent upon the history of previous shapes/poses [19]. They are also known to change their shape depending on temperature and that the temperature of the mirror depends on the electric current flowing through the actuator coils [20]. Both effects complicate their use for high speed refocusing devices.

In this paper, we present an image-based optimisation approach using an Alpao DM to achieve refocusing at 26.3 refocus sweeps per second over a defocus range −50 to 50 μm. The refocusing and optimisation procedure was applied to both a 40×/0.85 NA air objective and also a 40×/0.80 NA water immersion objective. Through the imaging of a star-test mask we determined that the mean of the estimated Strehl metric across the field of view for the 40×/0.80 NA water immersion objective was >0.7 over a field of view (FOV) of 200×200 μm^2^ for a refocus range of 52 μm and >0.6 over a 200×200 μm^2^ FOV for a refocus range of 77 μm.

## 2.1 Methods

### 2.2 Optical setup

The optical setup used for DM optimisation and testing is shown in Figure 2. A pair of aspheric lenses (AL1&2, 350230, Geltech/Thorlabs) imaged the LED (625 nm, M625L3, Thorlabs) onto a Lambertian diffuser (flashed opal glass diffuser, 50 DO 50, Comar Optics) that was placed in contact with and behind a custom star-test mask (JD Photo Data). The star-test mask is a 1.6 mm thick glass slide with one surface coated in a chrome layer that has an array of 1 μm diameter circular pinholes spaced on a hexagonal lattice with a period of 20 μm, which was used as the object during the deformable mirror (DM) optimisation procedure. The axial position of the star-test mask was controlled using a motorized stage (ESP100, Newport).

**Figure 2.**
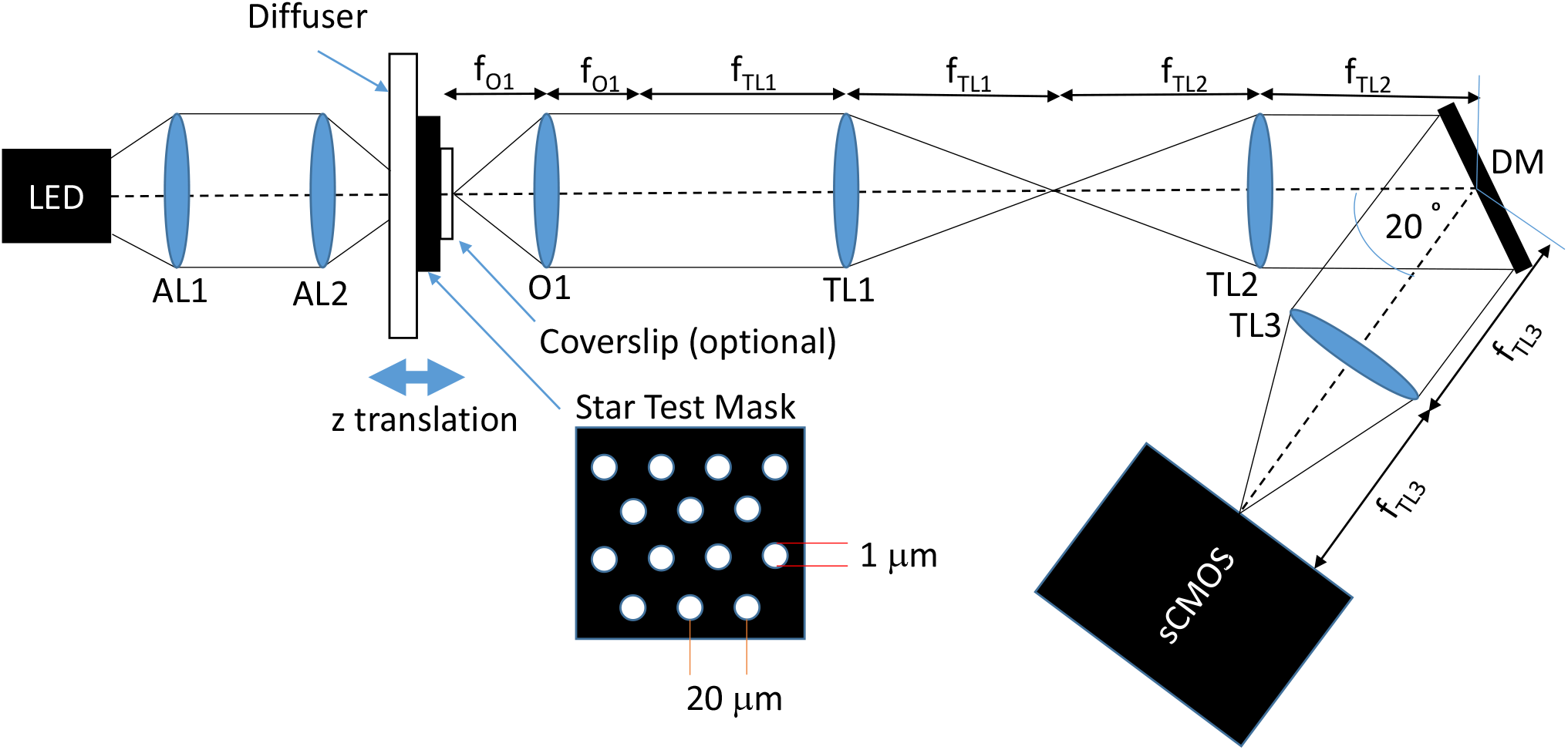
Optical setup used to optimise and test the deformable mirror for rapid refocusing. AL – aspheric lens; O – microscope objective, TL, tube lens; DM, deformable mirror.

Light transmitted through the star-test mask was collected by objective O1, which was either a 40×/0.85 NA air objective (1-UB827, Olympus) or a 40×/0.80 NA water-immersion objective (LUMPLFLN, Olympus). For the water immersion lens, as the system was aligned in a horizontal plane, we used ultrasound gel (UGEL250, Ana Wiz Ltd) as the immersion medium (which has a refractive index within 1% of that of water). The exit pupil of O1 was relayed via 4-f tube lenses TL1 (Thorlabs TTL100A) and TL2 (Thorlabs TTL200A), onto the deformable mirror (DM, DM97-15, Alpao), which applied a phase correction. Ideally, the pupil of O1 (diameter 7.65 mm and 7.20 mm for the air and water immersion objectives respectively) would be imaged onto the DM so that the pupil image matches the size of the DM (diameter 13.5 mm). However, in this setup the image of the pupil overfilled the DM, thereby reducing the NA of the system to 0.75.

In order to avoid using a beamsplitter, the DM was angled with its normal at 10º to the optical axis of O1 and TL1&2, and reflected the refocused wavefront to TL3 (f = 200 mm, TTL200A, Thorlabs), which produced an image on the sCMOS camera (ORCA-Flash4.0 v3, Hamamatsu) with an overall lateral magnification from the star-test mask of 22.2×. The camera was operated in its global reset edge-trigger mode with a 0.5 ms LED illumination pulse timed to occur when all rows of the sensor were exposed. To correct for the distortion introduced by the tilt of the DM with respect to the incident beam, the patterns applied to the DM were scaled by a factor of 1/cos(10º) in the direction parallel to the plane of the optical table.

### 2.2 Model describing DM surface profile

The optical path difference, OPD, required in the exit pupil of an objective lens to achieve a specific amount of defocus can be estimated for an on-axis defocused point by exploiting the sine condition, the expression for which will be referred to as high-NA defocus [7], and is,

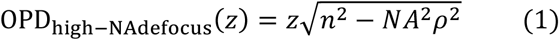

Here, *n* is the refractive index in the sample, *ρ* is the normalised radial pupil coordinate and *z* is the axial displacement of the object from the focal plane towards the objective. To account for aberrations of the objective lens and optical relay system, the dynamic response of the continuously oscillating DM as well as the potential need to under or over-drive the DM in order for it to reach the correct shape at the correct time, a series of Zernike modes were added to this to give a combined OPD for the mirror command signal,

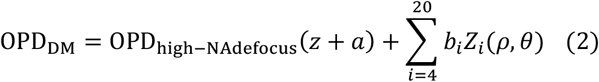

The Zernike modes *Z* are indexed by their Noll index *i* and each mode is normalised so that its inner product with itself over the unit circle is *π*. The amplitudes *a* and *b_i_* were determined by the optimisation procedure below.

### 2.3 Calculation of Strehl ratio and optimization metric

The Strehl ratio is defined as the ratio of the intensity at the centre of a system’s PSF to the theoretical maximum diffraction limited intensity that would be obtained in the absence of any aberration. A system with a Strehl ratio greater than 0.80 is commonly considered as being diffraction limited [21]. Figure 3(a) shows an example of a typical image acquired by the camera of the star-test mask using the 40×/0.8 NA air objective.

**Figure 3.**
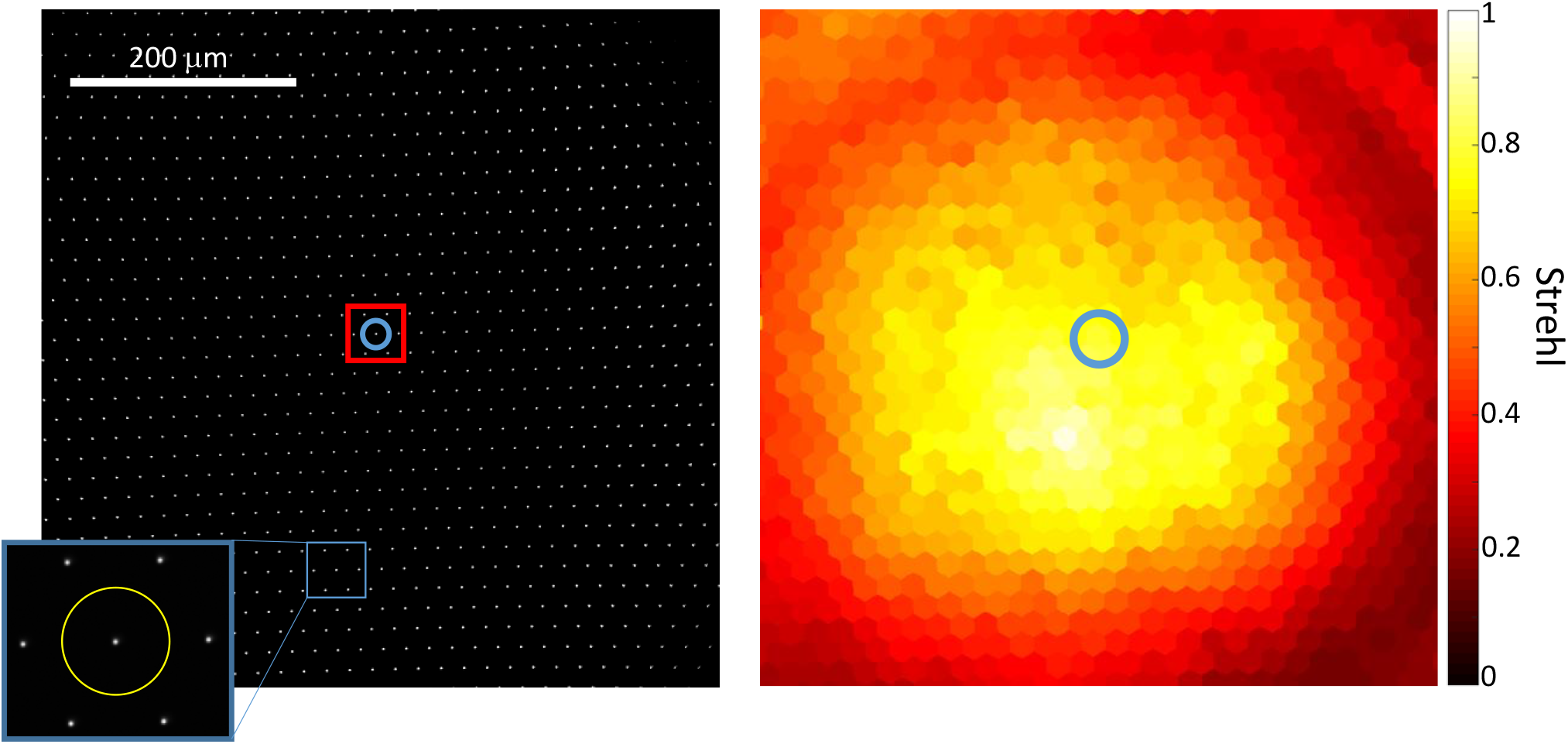
(a) Example raw image of the star-test mask (the contrast of the image has been increased to improve visibility of the pinhole images), recorded with the star-test mask in the focal plane of the 40×/0.85 NA air objective. The inset shows a zoomed-in region of the pinhole within the blue square. The yellow circle in the inset shows the circular bounding area around that pinhole. The perimeter of the yellow circle was used to find the local level of background light in the image for that pinhole and the values within the yellow circle were used to estimate the Strehl ratio. The red square of side length 60 μm shows the pinholes used to find the mean Strehl ratio, which is the score used in the optimisation. (b) False-colour Strehl map where each hexagon shows the Strehl metric calculated for a particular pinhole. The estimate for the Strehl ratio for the pinhole inside the blue circle in (a) is reported by the hexagon in the blue circle in (b).

The Strehl ratio was estimated for each pinhole using the following procedure. First, as the Strehl ratio is very sensitive to the background light in the image, we estimated the local background by taking the mean of pixel values lying at half the period of the star-test pattern from each pinhole, see yellow circle shown in inset to 3(a), and subtracted this from the image of that pinhole lying within the yellow circle. Next, the normalised maximum pixel value for each pinhole was found by dividing the maximum pixel value within each circle by the sum of all pixel values within the circle. A Strehl metric was then estimated by dividing the normalised maximum pixel value by the theoretical diffraction-limited value for the normalised maximum pixel value in the absence of any aberration, which was estimated using the theoretical Airy PSF convolved with the 1 μm diameter top hat function for the pinhole, and then allowing for the pixel size of the detector (6.5 μm). Finally, the Strehl metric for each pixel was displayed on a map (Voronoi diagram), where each hexagon corresponds to a specific pinhole; the Strehl metric for the pinhole marked with a blue circle in Figure 3(a) is shown by the false-colour of the hexagon identified with a blue circle in Figure 3(b).

These Strehl-metric maps show the spatial variation of Strehl metric across the field of view and were used to evaluate the performance of the DM when refocusing the optical system by different amounts.

Ideally, the DM should refocus the objective with an as large as possible diffraction-limited field of view at each defocus. We chose to use the mean Strehl value of the pinholes contained within a central 60 μm square of the camera field of view (shown in Figure 3(a) as a red square) as the metric for the optimisation algorithm.

### 2.4 DM performance

The Alpao DM97-15 is provided with a factory-measured influence matrix, and it is initially necessary to find the set of actuator commands that flatten the mirror, which was performed using a Shack-Hartmann wavefront sensor (HASO4VIS, Imagine Optics). To achieve this, light from a white-light source (Ocean Optics, Halogen HL-2000-FHSA) coupled via a 400 μm diameter multimode step-index fibre was collimated by an achromatic doublet (AC254-200-A-ML, Thorlabs), transmitted through a 50:50 non-polarising beam splitter and incident normal to the surface of the DM. After the DM, the reflection from the 50:50 non-polarising beamsplitter was directed through a 4× de-magnifying telescope, consisting of a pair of achromatic doublets spaced by the sum of their focal lengths AC254-200-A-ML, AC254-050-A-ML, Thorlabs), to create an image of the DM at the front focal plane of the Shack-Hartmann lenslet array. An iterative optimisation was then employed to flatten the DM based on the wavefront sensor measurement. The manufacturer-supplied influence matrix was used to calculate actuator commands for a particular desired surface and were added to the best flat commands in order to produce accurate representations of the desired surface. This was validated by commanding the DM to adopt a range of Zernike modes, and then confirmed using the Shack-Hartmann wavefront sensor.

The DM’s viscoelastic creep, which has a time constant of the order of 6 mins [19], and the heating effects of the actuator coil currents, which occur on a scale of 10s of seconds [20], mean that the mirror response depends on the previous actuator command history, which can be a significant issue.

Figure 4(a) shows the effect of visco-elastic creep. The mirror was either held in a continuous 50 μm high-NA defocus pose for 1000 s (blue curve) or oscillated from 50 μm to −50 μm high-NA defocus (each pose held for 200 ms) for 1000 s with the mirror on average flat (brown curve). Then mirror was then set to flat (*t =* 0 s) and the mean Strehl ratio of the central 200×200 μm^2^ of the image recorded for 1000 s. When the mirror was not on average flat for times before *t =* 0 s, then the mirror took longer to return to the initial flat pose, and hence the mean Strehl ratio took longer to return to its initial value, than when the mirror was oscillating. This shows that the unwanted effects of visco-elastic creep can be reduced by keeping the average pose on the mirror flat.

**Figure 4.**
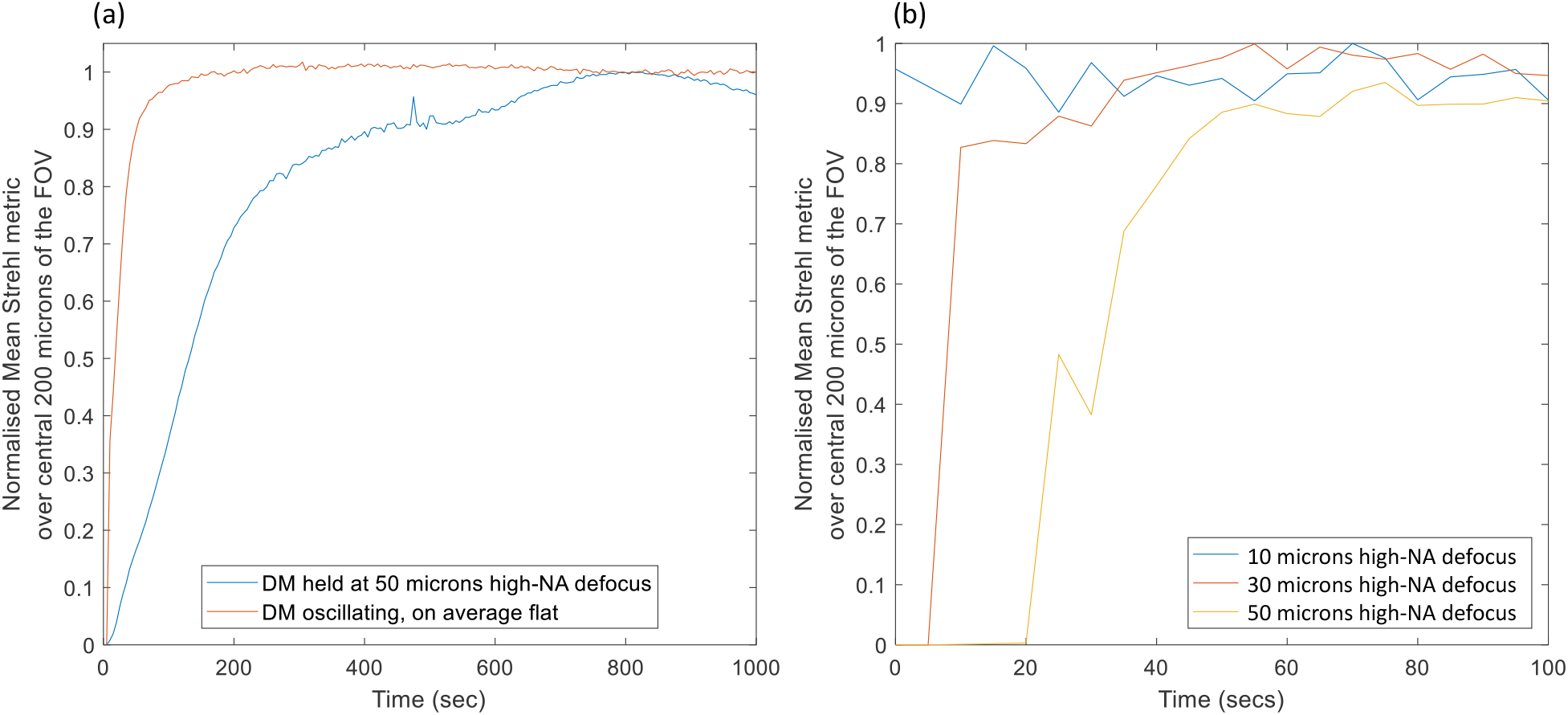
(a) Normalized mean Strehl ratio over the image of the central 200 μm of pinholes (normalized to the mean Strehl value obtained with a flattened DM), as the mirror returns to a flat pose after having been held in a position of 50 μm of high-NA defocus for 1000 s (blue) compared with the mirror oscillating from 50 μm to −50 μm of high-NA defocus for 1000 s (brown) with each pose held for 200 ms. The plots show that the visco-elastic creep can be avoided by keeping the average pose of the mirror flat, however there is still significant thermal creep. (b) Shows the same readout as for but where the mirror was oscillated with an average flat pose for different amounts of high-NA defocus for 120 s prior to measurement. Each pose was held for 100 ms during the oscillation. The plots show that the greater the amplitude of the oscillation then the greater the thermal creep and the longer it takes the mirror to return to flat.

Even when the mirror is on average flat, the mirror still takes some time to return to the initial flat pose. This was also allow any viscoelastic creep to occur if the mirror had been switched off prior to measurements. The dependence of the thermal creep on oscillation amplitude meant re-optimisation was necessary if the amplitude or frequency of the mirror oscillation was changed. attributed to the effect of thermal creep of the mirror surface, as the average drive current to the mirror actuators is higher when the mirror is oscillating compared to when it is flat. Figure 4(b) shows the effect of thermal creep of the mirror. The mirror was oscillated for 120 s through different levels of high-NA defocus, ranging from −10 to 10 μm to 50 to −50 μm, where each pose was held for 100 ms and where the mirror is always on average flat. The plots show the normalized mean Strehl ratio over the central 200×200 μm^2^ of the image as the mirror returned to a flat pose once the oscillation had ceased (normalized to the value of the mean Strehl when the mirror was flat). The greater the amplitude of the oscillation the more heat is generated by the mirror and so the longer it takes for the mirror to flatten.

We found (data not shown) that the viscoelastic and thermal creep prevented us from optimizing the mirror unless we waited at least 1 minute between each change of the mirror pose, which made the optimization of multiple mirror poses through a range of defocuses prohibitively slow (>12 hours) and also meant that the mirror was not able to oscillate rapidly through the range of poses.

Provided that the time-average shape of the mirror remained constant over a timescale shorter than the time constant of the creep, then the viscoelastic creep did not present a problem. We therefore chose to oscillate the DM continuously through a series of poses chosen so that the time-averaged DM pose remained flat – we found this to be necessary to successfully optimize the DM poses. The thermal creep could be minimised by allowing the mirror to oscillate through this series of poses for several minutes prior to any measurement in order to allow the system to reach thermal steady state. Figure 4(b) shows that at least 60 s was required for this. We chose to use a period of 10 minutes as this would also allow any viscoelastic creep to occur if the mirror had been switched off prior to measurements.

The dependence of the thermal creep on oscillation amplitude meant re-optimisation was necessary if the amplitude or frequency of the mirror oscillation was changed.

### 2.5 Optimisation algorithm

The approach used involved optimising a set of control mirror poses corresponding to a discrete set of defocus positions of the star-test mask (−50, −40,…, 40, 50 μm, positive defocus is defined here in the direction from the object towards the objective). For each defocus position, first the motorized translation stage was used to move the star-test mask to the required defocus position. A range of high-NA defocus poses were then applied to the DM centred on the required defocus position and the Strehl metric measured for each one. The amount of high-NA defocus providing the highest Strehl metric was then determined from a quadratic fit to the data points spanning the maximum and 3 points either side of the maximum, see Figure 5. This value was then applied to the DM and the Strehl metric found. If this was greater than the previous best metric, then this was taken to be the new best pose, otherwise the previous value was retained – we found that this reduced the chances of noise affecting the convergence of the optimisation. The above procedure was then carried out for Zernike modes with Noll indices from 4 to 20 in turn. The whole optimisation process was then repeated until there was no further change, which typically required 4 cycles.

**Figure 5.**
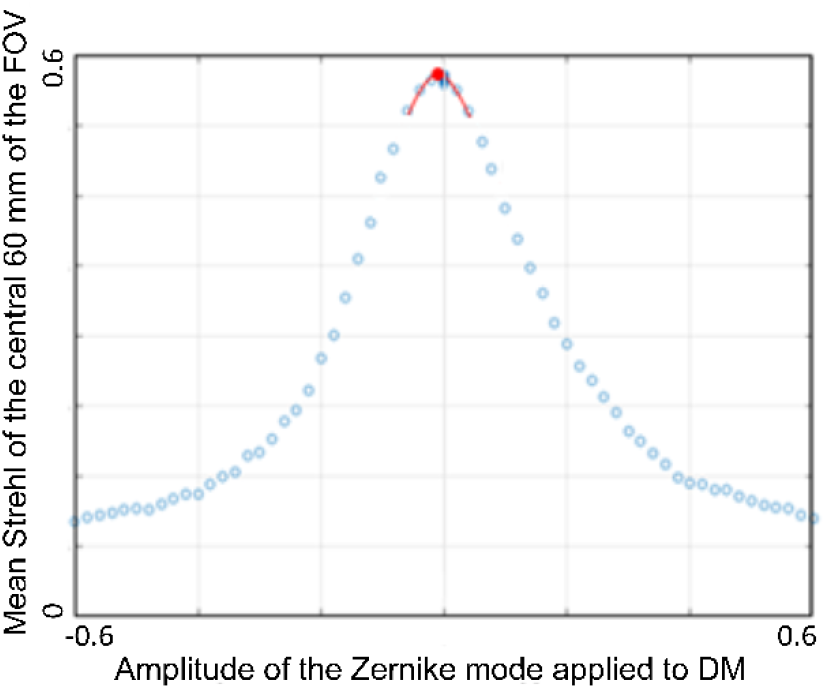
Plot of the mean Strehl metric as a function of the amplitude of the Zernike mode for Noll index 4. The red line shows estimation of the location of the maximum found by fitting a quadratic to the data points spanning the maximum and 3 points either side of the maximum.

## 3. Results and discussion

### 3.1 Static optimisation

The DM was optimized initially for each amount of desired defocus in turn without continuously oscillating through a sequence of poses. The optimisation approach described in section 2.5 was modified, adding a 1 min pause after each change in mirror pose, to allow for most of the visco-elastic creep to occur, and to then apply the reverse profile to the mirror for 1 min, in order to keep the long-term average pose flat. This made the optimisation procedure very time consuming (>12 hours) and changes in ambient conditions such as temperature could cause the optimisation to fail. Results obtained for the Olympus 40×/0.85 air objective are shown in Figure 6. It can be seen that the largest field of view is achieved for zero refocus. The decrease in the field of view is more pronounced for positive defocus. The maximum and mean Strehl values over the central 200×200 μm square of the field of view as a function of defocus are shown in Figure 7. We achieved a maximum Strehl of >0.8 over a range of 59 μm, better than would be expected correcting only for primary defocus (Table 1). As well as applying high-NA defocus, the optimisation procedure corrected for system aberrations. The final performance was limited by field-dependent aberrations, in particular field curvature; the DM could only provide an average correction over the field of view for these kinds of aberrations.

**Table 1.**
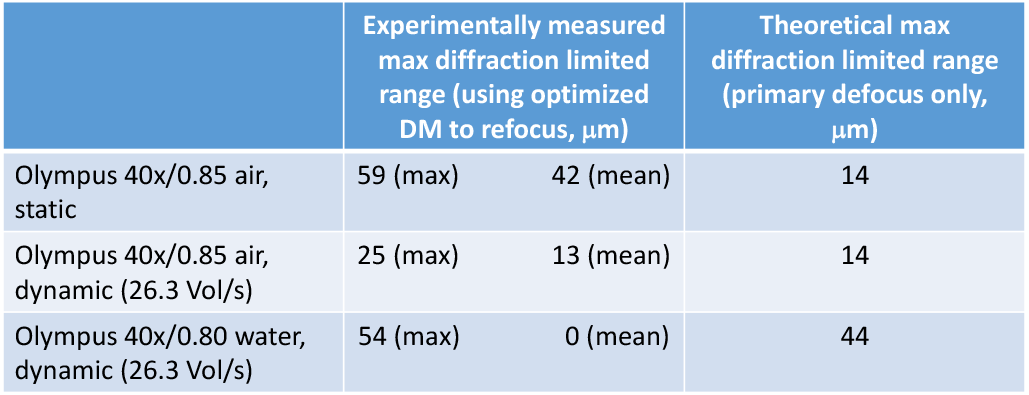
Summary of the diffraction-limited refocus ranges achieved experimentally using the DM – in terms of maximum Strehl at each defocus (max) and also the mean Strehl over the central 200×200 μm square of the field of view (mean) – and the equivalent maximum theoretical values for application of primary defocus alone.

| | Experimentally measured max diffraction limited range (using optimized DM to refocus, $\mu\text{m}$) | | Theoretical max diffraction limited range (primary defocus only, $\mu\text{m}$) |
| --- | --- | --- | --- |
| Olympus 40x/0.85 air, static | 59 (max) | 42 (mean) | 14 |
| Olympus 40x/0.85 air, dynamic (26.3 Vol/s) | 25 (max) | 13 (mean) | 14 |
| Olympus 40x/0.80 water, dynamic (26.3 Vol/s) | 54 (max) | 0 (mean) | 44 |

**Figure 6.**
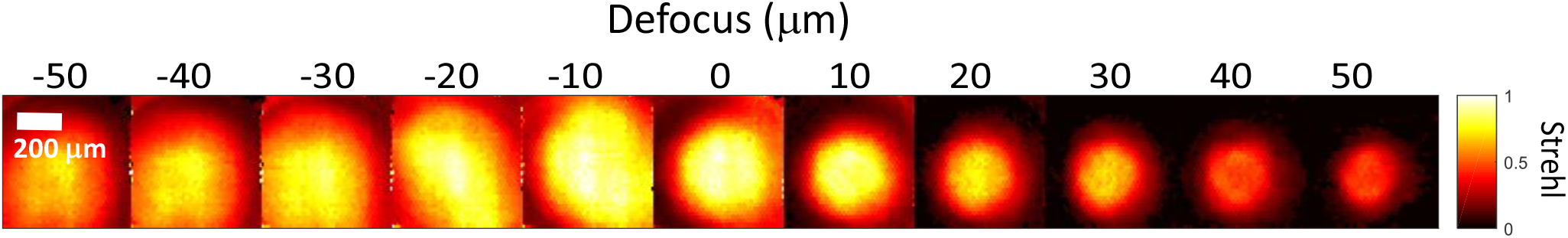
Static optimisation of the DM for the Olympus 40x/0.85 air objective. The results show the Strehl maps across the field of view (600×600 μm) for defocus positions −50 through to 50 μm.

**Figure 7.**
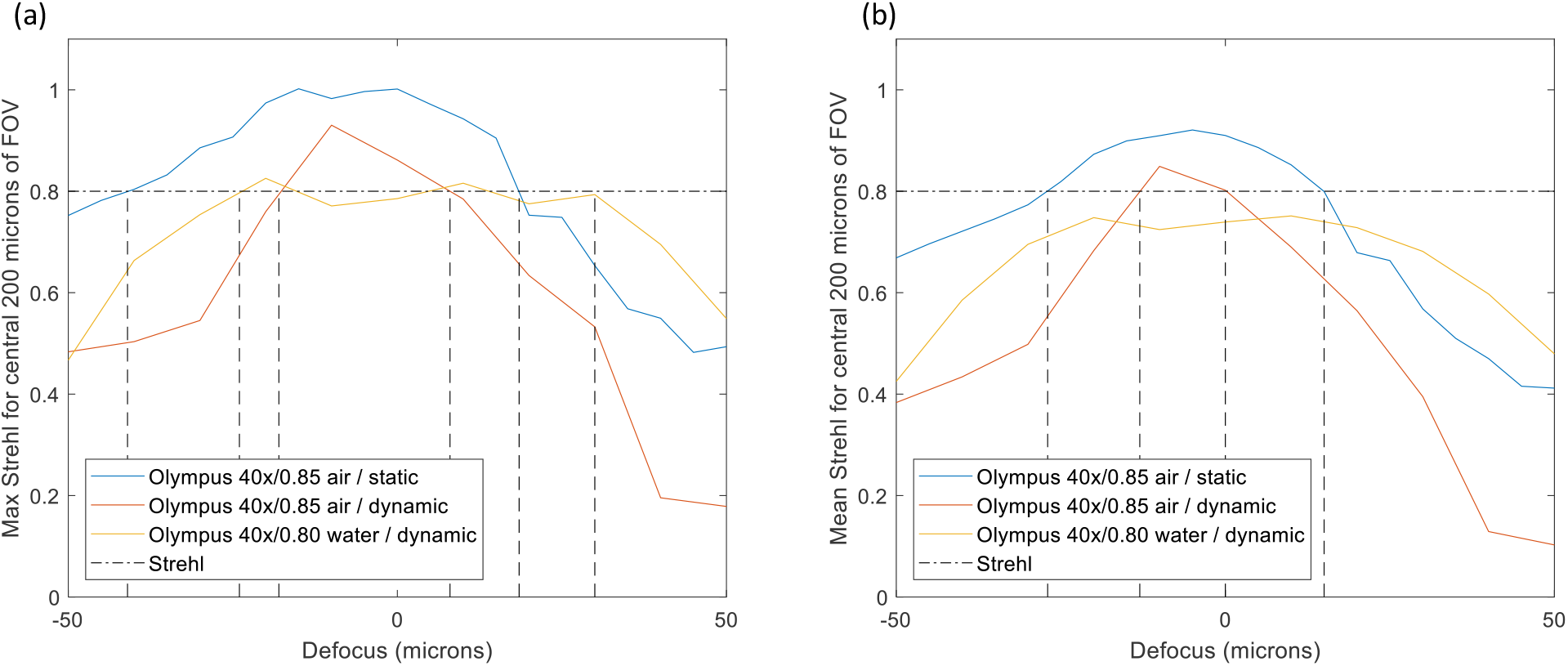
Maximum Strehl (a) and mean Strehl (b) of the central 200×200 μm square of the field of view for the statically optimised Olympus 40x/0.85 air objective, as well as the dynamically optimised (26.3 refocus sweeps/sec) Olympus 40x/0.85 air objective and Olympus 40x/0.80W water immersion objective. The dashed line shows the diffraction limit.

### 3.2 Dynamic optimisation

In order to achieve rapid refocusing, a periodic sequence of poses was sent to the mirror at its internal update period of 65 μs. We chose to optimise the outward sweep using a sequence of 11 control poses, corresponding to defocus positions of −50 to 50 μm in steps of 10 μm. For the return sweep, we chose to use 4 (non-optimised) poses of high-NA defocus in steps of 20 μm. We then used linear interpolation of the actuator commands between each consecutive control or calculated pose to generate 38 intermediate poses in order to achieve a smooth motion of the mirror surface. This resulted in a defocus sweep over the range −50 to 50 μm – including return to the start position – every 38.0 ms, i.e. at a sweep rate of 26.3 Hz. The total number of poses per sweep was 585.

To avoid viscoelastic creep, an offset was applied to the calculated return poses to ensure that the temporally averaged pose of the DM over the whole cycle remained flat. The control software (MATLAB), was implemented to ensure that the DM was continually oscillating at 26.3 Hz throughout the entirety of the optimisation procedure and afterwards in order to ensure that thermal creep effects settled out and remained constant. The initial 10 minute warm-up used unoptimized control poses for the calculated high-NA defocus. During optimisation, the illumination LED was synchronised with the mirror poses so that images of the star test mask were only acquired when the mirror had the pose corresponding to the control pose that was currently being optimised. This strobing method allowed the optimisation of each control pose separately. The whole optimisation procedure took approximately 90 minutes and re-optimisation, including re-flattening of the DM, was required on a daily basis, which was attributed to variations in ambient temperature.

Following the optimisation, there was an evaluation step where images of the star-test mask were obtained for each defocus position with the star-test mask displaced (Δ*z*) by −4 to 4 μm from the position used during optimisation, and Strehl maps obtained, see columns in Figure 8(a) for the results for an Olympus 40x/0.85 air objective. The data from each defocus position were then combined to produce the best Strehl map for each optimised defocus position (Figure 8(b)). The final column (Figure 8(c)) shows the Δ*z* value at which the best Strehl value was obtained and therefore indicates the amount of field curvature present at each optimised defocus position. The field curvature increased with defocus and reversed in sign through the focal plane. Figure 9 shows results for the 40×/0.80W water dipping objective – where ultrasound gel was used as the immersion medium. The results also show some field curvature. Figure 7 shows a plot of (a) the maximum, and (b) the mean Strehl metric over the central 200×200 μm^2^ of the camera FOV for both objectives as a function of optimised defocus position for both static and dynamic optimisations. For the water dipping objective, a mean Strehl of >0.6 is obtained over a defocus range of 80 μm; for the air objective this range is 45 μm. For the 40x/0.85 air objective, the performance achieved with the dynamic optimisation is lower than that achieved with the static optimisation, which we attribute to the motion of the DM during the 0.5 ms LED illumination during both optimisation and evaluation measurements (during the LED illumination the DM refocuses a distance of 2 μm at 26.3 volumes/s).

**Figure 8.**
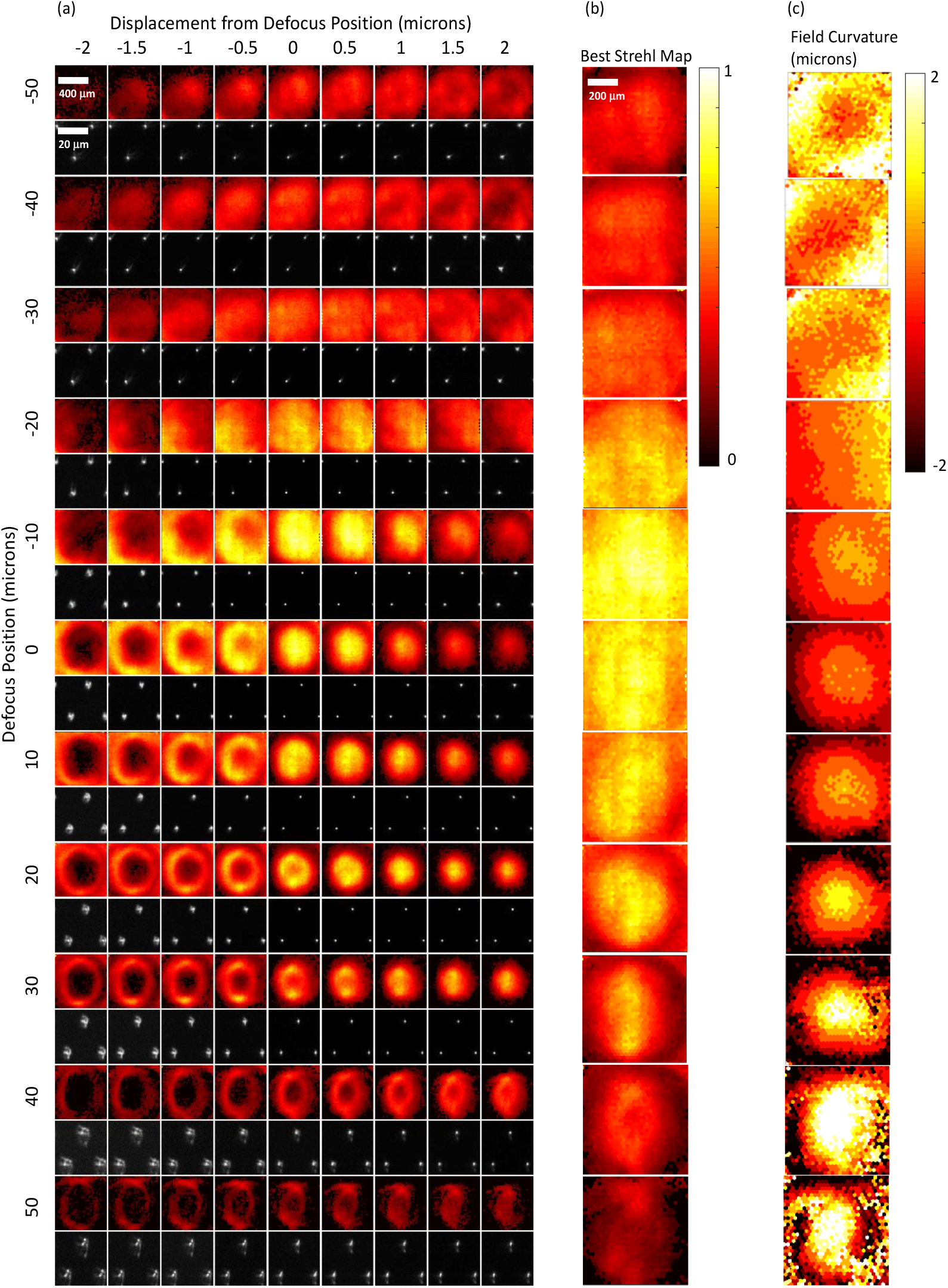
Results obtained from dynamic optimisation of the DM at 26.3 Hz with the 40×/0.85 air objective and coverslip. a) False-colour Strehl maps (upper row) and small ROI of raw star test mask image from the centre of the camera’s FOV (lower row) for DM poses optimised to provide defocuses of −50 to 50 μm in 10 μm steps (top to bottom). For clarity, the brightness of each raw star-test mask image has been individually scaled to the maximum for that image. For each optimised DM pose, data is shown for star-test mask defocus positions Δ*z* of −2 to 2 μm in 0.5 μm steps away from the defocus position used during DM optimisation (left to right). b) False-colour map of best Strehl value for each pinhole taken over all Δ*z* values. c) False-colour map showing Δ*z* location in units of μm of the best Strehl value for each pinhole, i.e. showing the curvature of the field imaged.

**Figure 9.**
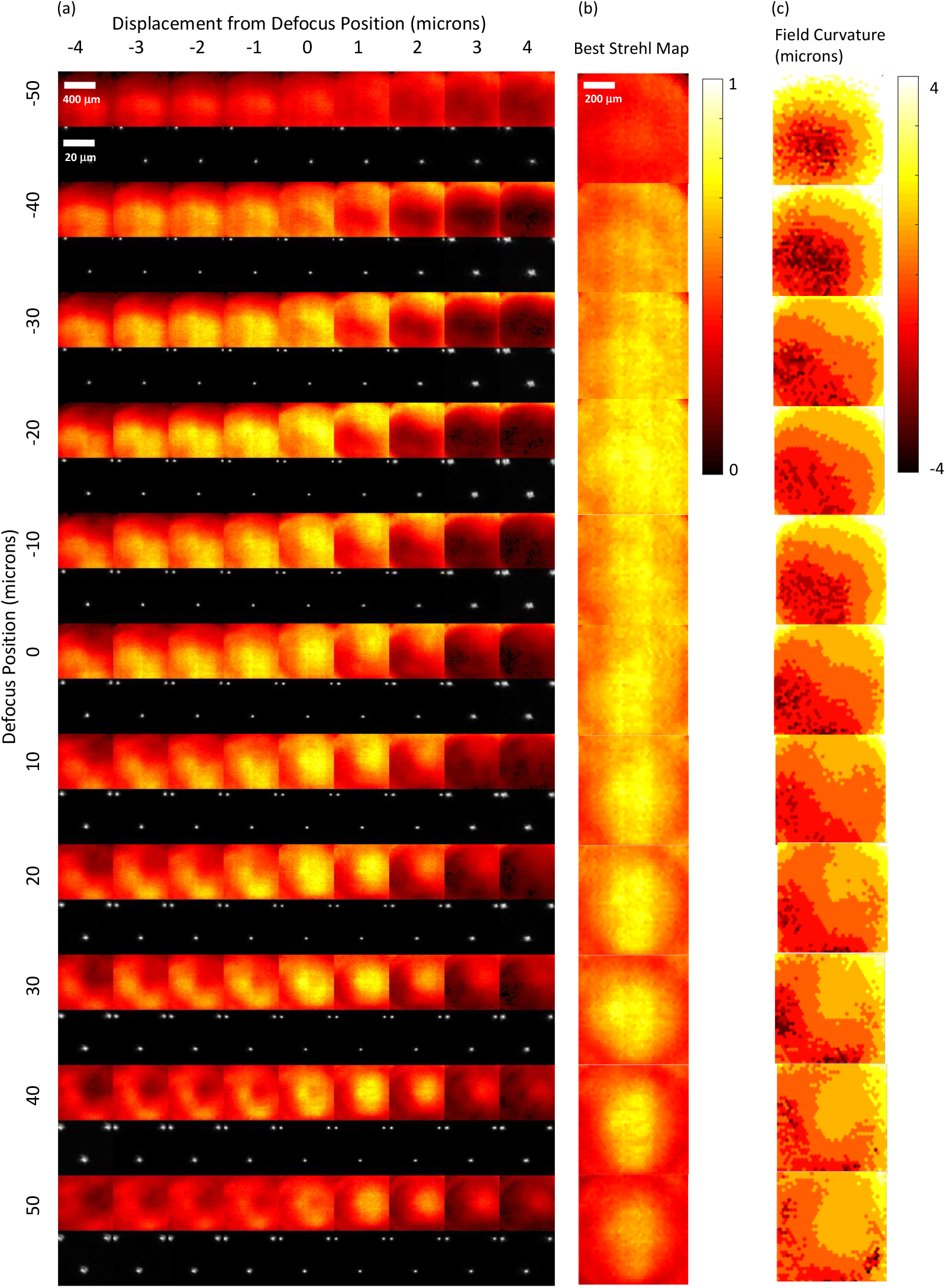
Results for dynamic optimisation of the DM at 26.3 Hz for the 40×/0.80 water dipping objective with ultrasound gel as the immersion medium. a) False-colour Strehl maps (upper row) and small ROI of raw star-test mask image from the centre of the camera’s FOV (lower row) for DM poses optimised to provide defocuses of −50 to 50 μm in 10 μm steps (top to bottom). For clarity, the brightness of each raw star-test mask image has been individually scaled to the maximum for that image. For each optimised DM pose, data is shown for star-test mask defocus positions Δ*z* of −4 to 4 μm in 1 μm steps away from the defocus position used during optimisation (left to right). b) False-colour map of best Strehl value for each pinhole taken over all Δ*z* values. c) False-colour map showing Δ*z* location in units of μm of the best Strehl value for each pinhole, i.e. showing the curvature of the field imaged.

For both the air and water dipping objectives in the dynamic optimisation, the amount of each mode applied for each defocus position is shown in Figure 10. Beyond *Z*_13_, additional Zernike modes provided no further improvement in the mean Strehl metric. The dashed horizontal lines in this figure show the amount of each mode alone that would be required to exceed the diffraction-limit RMS error in order to give an indication of the scale of the correction applied.

**Figure 10.**
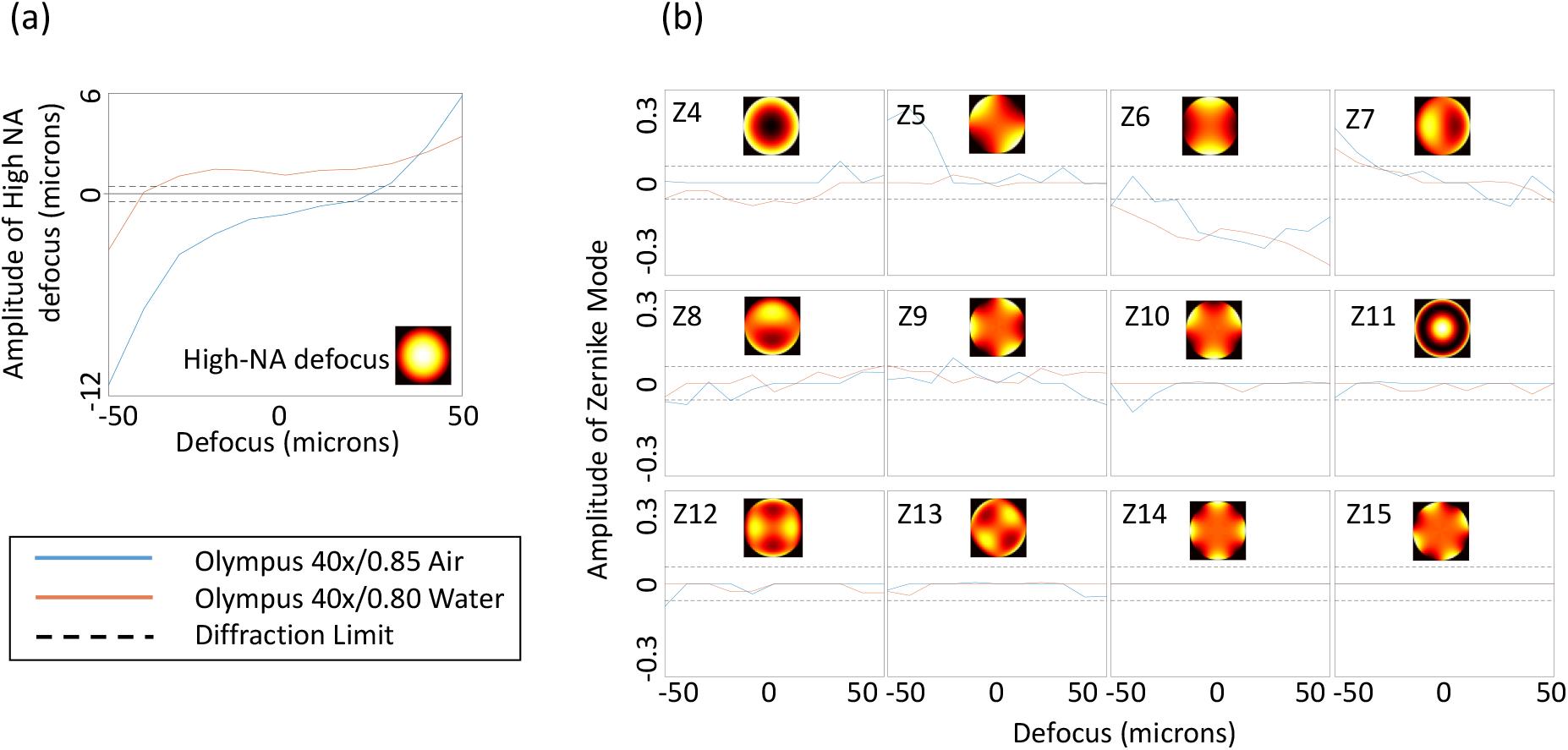
Plots showing the amplitude of the high NA defocus mode and Zernike modes (indexed by Noll index), for each defocus. Blue curve shows results for the 40×/0.85 lens with coverslip. Red curves show results for the 40×/0.8 water lens with ultrasound gel as the immersion medium. The black dashed lines show the amplitude above which the RMS wavefront aberration for that mode alone exceeds the diffraction limit of *λ*/14.

To further validate the performance of the system for the 40×/0.80W objective, the DM was set to sweep at 26.3 Hz through the set of dynamically optimised poses with the camera acquiring an image of the star-test mask (with a reduced FOV) for every optimised DM pose, i.e., 11 images per oscillation of the mirror. During this process, the star-test mask was moved with a constant axial velocity from −50 to 50 μm at a rate of 6 μm s^−1^. Figure 11 shows a montage of the data acquired, with the images from each sweep (defocus position) shown as a column and with data from every 30^th^ sweep shown moving from left to right. It can be seen that the images where the star-test mask is in focus (diagonal) tracks the motion of the stage and validates the DM defocusing at video frame rate.

**Figure 11.**
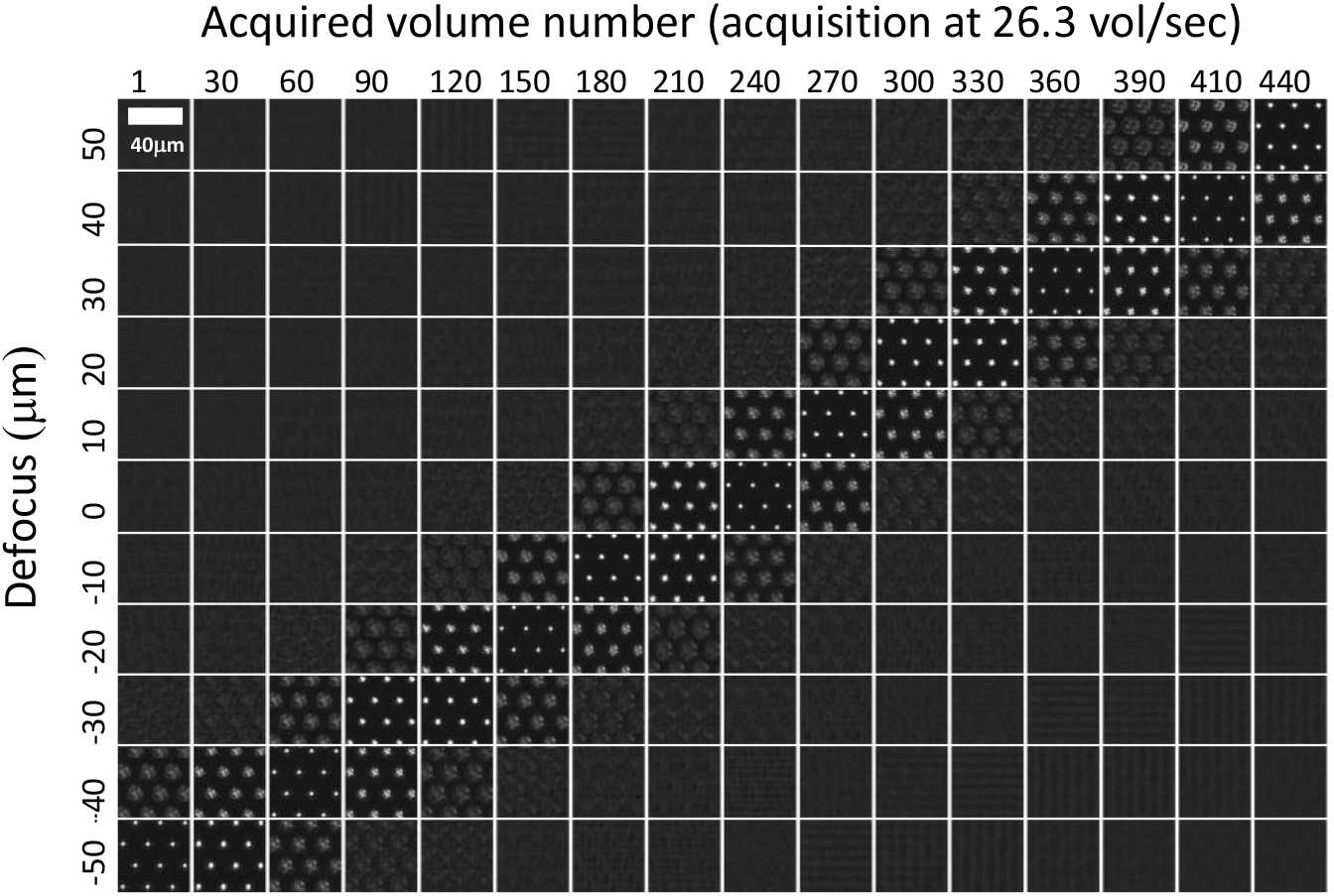
Image sequence acquired from every defocus position as a function of time as the star-test mask is translated at a constant velocity of 6 μm s^−1^ towards the microscope objective. Images were acquired using the 40×/0.80W objective with ultrasound gel as the immersion medium. Refocus sweeps were performed at 26.3 Hz. Each column shows a sequential set of sCMOS images acquired at each of the 11 control poses of the DM, which correspond to refocus positions over the range −50 to 50 μm. Every 30th DM sweep/volume is shown moving left to right across the figure, with the number of the sweep/volume acquired shown at the top of each column.

## 4. Conclusions

We have investigated the use of an Alpao DM97-15 deformable mirror to perform rapid remote refocussing of a microscope at a rate of 26.3 Hz through a defocus range of − 50 to 50 μm. The mirror was optimised for 11 control poses per sweep, with 4 high-NA defocus plus offset poses being used for the return sweep and with linear interpolation to generate intermediate poses. The entire optimisation procedure was conducted with the mirror oscillating continuously at the desired sweep rate in order that the temperature of the DM remained constant and so to avoid issues with thermal creep of the DM. Visco-elastic creep was avoided by keeping the temporal average profile of the mirror constant.

The performance of the system was tested using a 40×/0.85 air objective and a 40×/0.80 water objective. The air objective enabled a mean Strehl metric of >0.6 over a field of view of 200×200 μm^2^ and for a refocus range of 40 μm to be achieved. The water objective with ultrasound gel immersion fluid achieved a mean Strehl metric of >0.6 over 200×200 μm^2^ over a larger refocus range of 77 μm.

The results showed increasing field curvature for increasing defocus and showed that there is a limit to the amount of defocus that can be corrected with the deformable mirror for the configuration used here. The DM in our setup is unable to correct for field dependent aberrations such as field curvature in its location conjugate to the pupil of the objective. The refocus rate of the DM was limited by the power of the LED light source used during optimisation and much faster sweep rates should be achievable with a brighter source.

We plan to apply this DM refocusing system in a LSFM where the illumination light sheet position is swept along the optical axis of the detection objective in synchrony with the refocusing provided by the DM in order to achieve video-rate volumetric LSFM with high fluorescence collection efficiency and further, to use the DM to correct for previously-determined/estimated depth-dependent sample-induced aberrations.

## Acknowledgements

The authors gratefully acknowledge funding from the British Heart Foundation (BHF, NH/16/1/32447). TW acknowledges a PhD studentship funded by the Engineering and Physical Sciences Research Council. The authors declare no conflicts of interest.

## Appendix

The following expressions were derived using Mathematica (Wolfram).

The amount of primary defocus (corresponding to an axial displacement *z*_1_) that best corrects for high-NA defocus of *z*, in terms of minimising the RMS difference between applied primary defocus and desired high-NA defocus over the pupil of the objective is,

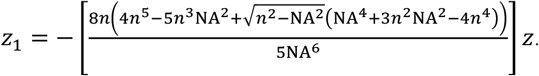

Here, *n* is the refractive index of the immersion medium, λ is the wavelength of light in vacuum and NA is the numerical aperture.

The standard deviation (StdDev) of the difference between the optimal primary defocus and the high-NA defocus at a particular defocus *z* is,
StdDev(Z) =

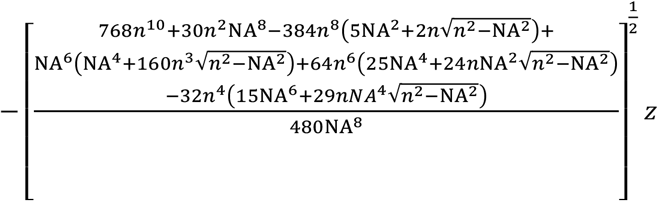

The maximum defocus that can be achieved before the
optimum amount of primary defocus (*z*_1_) no longer provides a diffraction-limited approximation to the high-NA defocus,

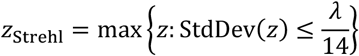

is given by,

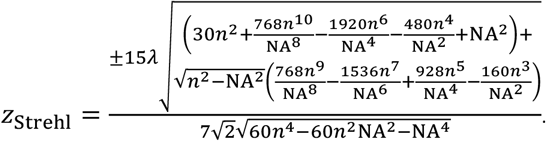

## Notes

### Competing Interest Statement

The authors have declared no competing interest.

